## Supplemental S1 Fig for "Participatory science methods to monitor water quality and ground truth remote sensing of the Chesapeake Bay"

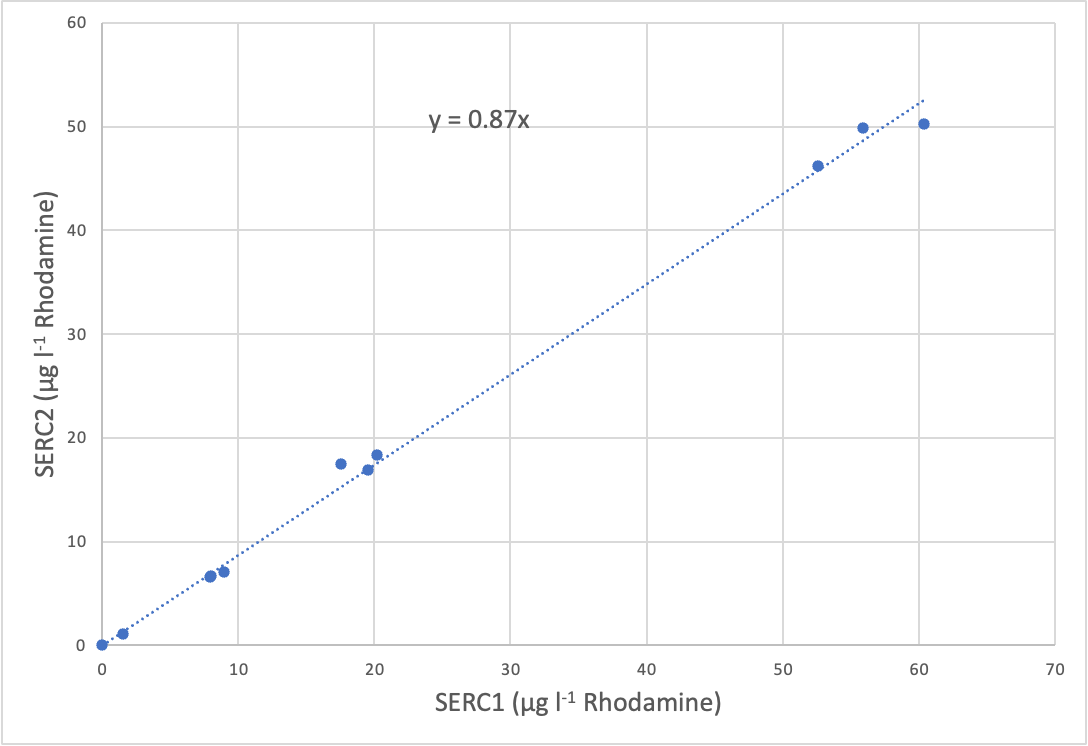

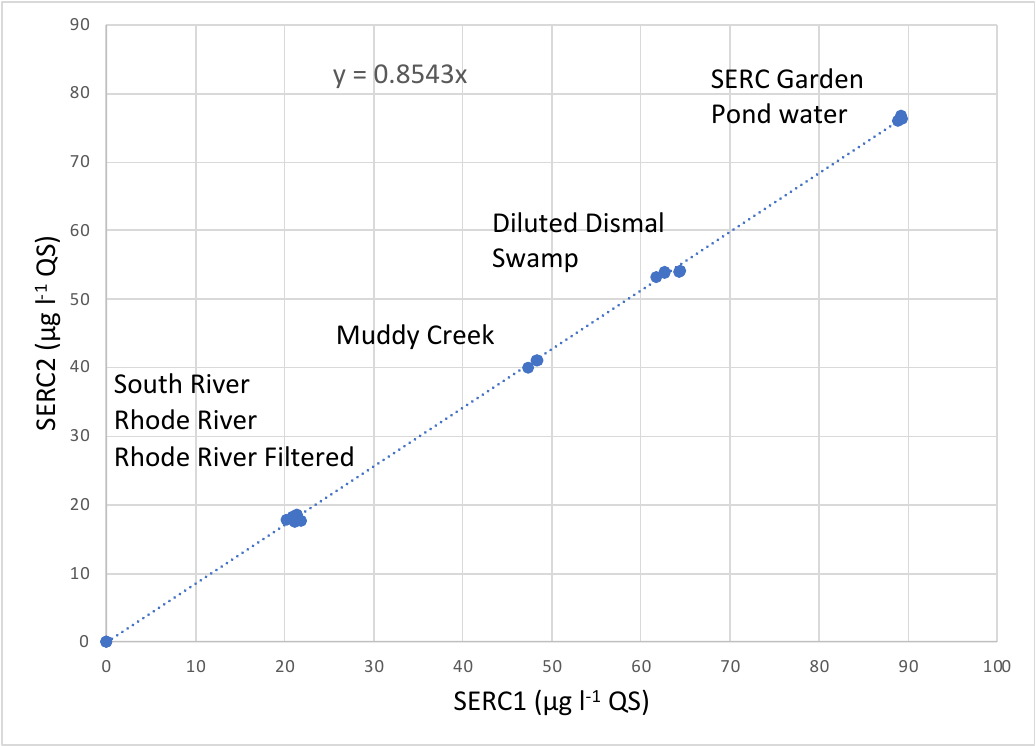
**S1 Fig Multiple sample comparison of Aquafluor units SERC1 and SERC2 CDOM and IVChl measurements.** Samples from multiple environments of varying salinity and spanning a range of CDOM (a) and in vivo chlorophyll-a fluorescence (IVChl) (b) content were compared to derive an overall proportion between the two instruments. All SERC2 CDOM and IVChl values in this report are the Fieldscope values divided by 0.85 and 0.87, respectively. “Rhode River Filtered” refers to a sample that has been filtered using the CDOM protocol to remove phytoplankton. The graph shows that filtration did not change the CDOM reading compared to using a whole sample. Line shows linear regression with zero intercept assumed, fitted equation is annotated.

b

a
