## Supplemental S1 File for "Participatory science methods to monitor water quality and ground truth remote sensing of the Chesapeake Bay"

**S1 File. Developer Testing and User experience with HydroColor v2**

The new RAW data image processing was tested using an exposure target that contains 25 different reflectances ranging from 4.5 to 72%. Each square of the exposure target was captured in a separate picture under constant illumination. The camera measured light levels were normalized to the 18% reflectance square (the same as it is done in a traditional HydroColor measurement). The camera measured reflectance was compared to the stated reflectance of each square. Data from three test smartphone cameras, an iPhone SE, Samsung Galaxy S21 and Google Pixel 5, showed that target reflectance was measured consistently among all the smartphones (Figure 1). Error in camera measured reflectance was < 1% relative to the reflectance standard for all cameras, based on the regression between camera measurements and stated standard reflectance. Although this specific test was not performed with HydroColor V1, intercomparison of compressed image data and calibrated radiometer showed some nonlinearity in the response (Leeuw and Boss, 2018). In addition, the extent and form of the nonlinearity is expected to differ between phones, although no tests were performed to establish the extent of the differences.

In general, participants found the v2 app was easy to use. The most common challenge to completing an acquisition was unstable operation of the smartphone compass making it difficult to position to the correct azimuth. This appeared to occur most often in phones with Android operating systems. Re-calibrating the compass using the established “Figure-8” maneuver usually improved the response. Also, it was noted that the physical azimuth positioning differed slightly (~5 degrees) between two phones being deployed side-by-side, both supposedly at the correct azimuth. This is difference is within the normal accuracy of the smartphones (Allmendinger, et al. 2017) and the error in azimuth should have little effect on the estimate of
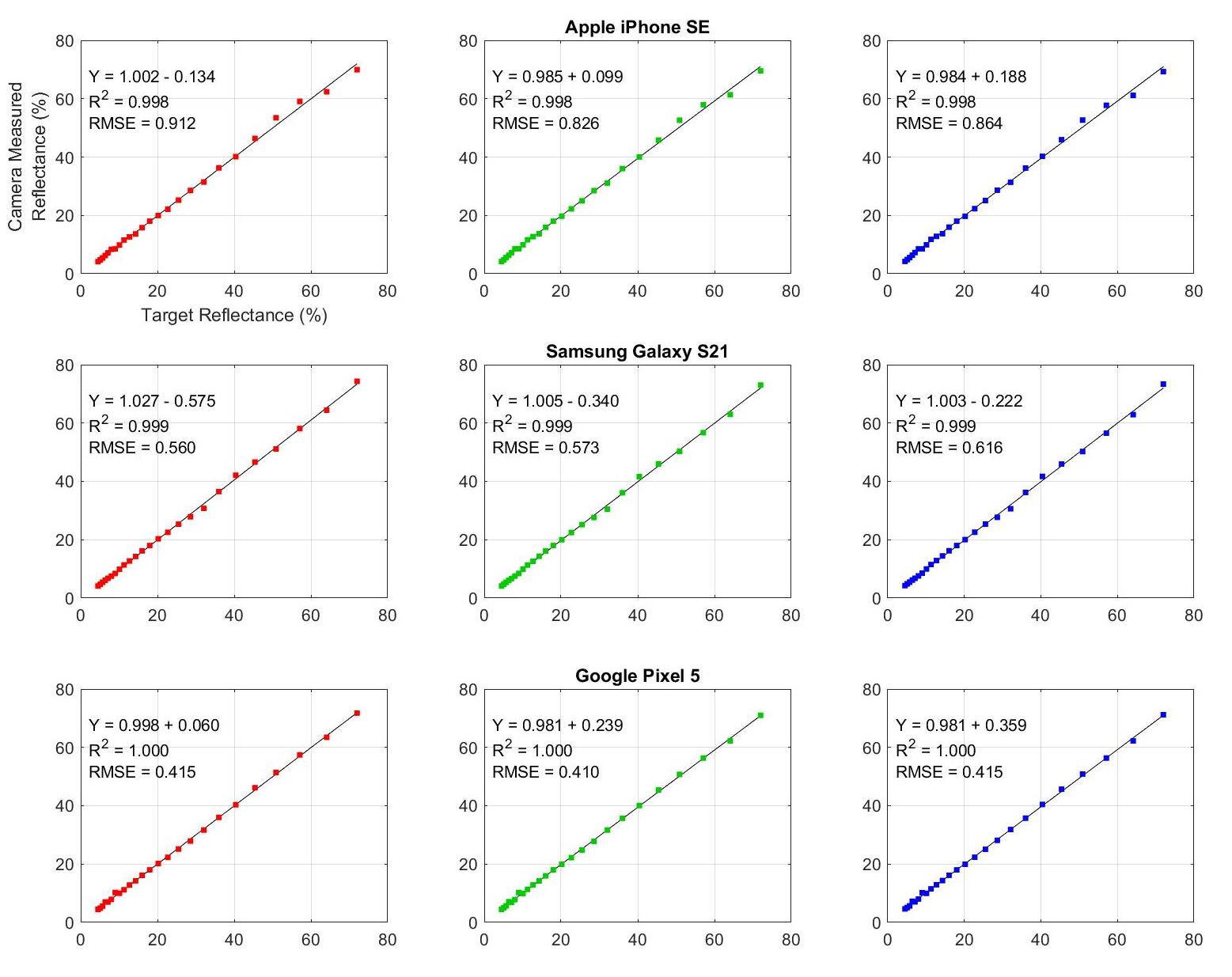
Rrs which in any case assumes a fixed sea surface reflectance (Leeuw and Boss, 2018).

**Fig 1 Verification of camera reflectance measurement using Sekonic exposure target.** 25 reflectance squares of varying reflectance were imaged with each device. The camera computed reflectance in each color channel is plotted against the specified reflectance of the exposure target square. Each row in the figure shows data from a specific device. The red, green, and blue color channels are shown in each column. Graphic provided by Thomas Leeuw.

References:

Allmendinger, R. W., Siron, C. R., & Scott, C. P. (2017). Structural data collection with mobile devices: Accuracy, redundancy, and best practices. Journal of Structural Geology, 102, 98-112. <https://doi.org/10.1016/j.jsg.2017.07.011>

Leeuw, T., & Boss, E. (2018). The HydroColor App: Above Water Measurements of Remote Sensing Reflectance and Turbidity Using a Smartphone Camera. Sensors (Basel, Switzerland), 18(1), 256. <https://doi.org/10.3390/s18010256>
