## Supplemental S1 Table for "Participatory science methods to monitor water quality and ground truth remote sensing of the Chesapeake Bay"

**S1 Table– Listing of the data entry types on the CWW Fieldscope site, serc.fieldscope.org.**

| **Field Name** | **Group** | **Units** | **Data Type** | **Categorical List** |
| --- | --- | --- | --- | --- |
| Date | Environmental Conditions |  |  |  |
| Time | Environmental Conditions | AM, PM |  |  |
| What was the percent cloud cover when collecting? | Environmental Conditions |  | categorical | Clear (0-5%), Mostly clear 25%, Partly cloudy 50%, Mostly cloudy 75%, Cloudy (90-100%) |
| What was the wind like? | Environmental Conditions |  | categorical | Calm, Light to Breezy, Breezy, Windy, Strong Winds |
| How were the waves? | Environmental Conditions |  | categorical | Completely calm, Rippled to small wavelets, Few whitecaps, Frequent whitecaps, Many whitecaps |
| Have you noticed submerged aquatic vegetation (SAV), also known as underwater grasses, growing in the surrounding area? | Submerged Aquatic Vegetation (SAV) |  | categorical | Yes, No |
| HydroColor Turbidity - 1 | HydroColor App | NTU | numeric |  |
| HydroColor Turbidity - 2 | HydroColor App | NTU | numeric |  |
| HydroColor Turbidity - 3 | HydroColor App | NTU | numeric |  |
| Type of phone | HydroColor App |  | categorical | Apple, Android |
| What was the Secchi depth? | Secchi Disk | meters | numeric |  |
| Secchi measurement taken on the shady side of the boat? | Secchi Disk |  | categorical | Yes - recording from the shady side of boat/ platform (preferred), No - sun directly overhead/ no shady side available |
| What was the total water depth? (if known) | Secchi Disk | meters or feet | numeric |  |
| Did you collect a water sample? | Water sample |  | categorical | No - please skip to Notes, Yes - I filled a sample tube, Yes - I filled a sample tube and validation bottle |
| Who analyzed the water sample? | Water sample |  | categorical | I analyzed the water sample, I sent the sample back to SERC, Someone from my local Riverkeeper organization analyzed the sample |
| Is there anything else you would like us to know? | Notes |  | open text |  |
| Instrument ID | Instrument Reading |  |  | SERC1-SERC8 |
| In Vivo Chlorophyll 1 | Instrument Reading | ug/L | numeric |  |
| In Vivo Chlorophyll 2 | Instrument Reading | ug/L | numeric |  |
| In Vivo Chlorophyll 3 | Instrument Reading | ug/L | numeric |  |
| CDOM 1 | Instrument Reading | ug/L | numeric |  |
| CDOM 2 | Instrument Reading | ug/L | numeric |  |
| CDOM 3 | Instrument Reading | ug/L | numeric |  |
| Turbidity 1 | Instrument Reading | NTU | numeric |  |
| Turbidity 2 | Instrument Reading | NTU | numeric |  |
| Turbidity 3 | Instrument Reading | NTU | numeric |  |

For categorical variables, the user selectable responses are listed.
