## Supplemental S2 Table for "Participatory science methods to monitor water quality and ground truth remote sensing of the Chesapeake Bay"

**S2 Table. Intercomparison of in vivo chlorophyll-a (IVChl) and CDOM fluorescence readings for six identically calibrated Aquafluor units.**

| **Aquafluor ID** | **Relative IVChl** | **Relative CDOM** |
| --- | --- | --- |
| **1** | 1.00±0.06 | 1.00±0.01 |
| **2^a^** | 1.02±0.01 | 1.00±0.01 |
| **3** | 1.03±0.03 | 0.96±0.01 |
| **4** | 1.10±0.03 | 0.98±0.01 |
| **7** | 1.01±0.02 | 0.94±0.01 |
| **8** | 1.07±0.02 | 0.95±0.01 |
| **Average Response Relative to SERC1** | 1.05 | 0.97 |

Post calibration readings using a sample from the Rhode River, Chesapeake Bay. Tabulated is the average (±SD) of three readings relative to the reference unit SERC1. To correct for drift in IVCHL reading, samples were read on unit 1 at start and end of the trial and the average reading was used for comparison.

^a^Instrument SERC2 uses LEDs with slightly different specifications than other units, separate tests showed that despite using identical calibration protocols these differences result in sample readings 15% lower than other units (see S1 Fig), the listed ratio is adjusted for this difference.
