## Supplemental S3 Table for "Participatory science methods to monitor water quality and ground truth remote sensing of the Chesapeake Bay"

**S3 Table. Results of multiple linear regression calculations with extracted Chl as the dependent variable and selected volunteer measurements as the independent variables.**

| **# of Terms** | **Intercept** | **IVChl** | **Turbidity** | **CDOM** | **Adjusted R^2^** | **Degrees of freedom** |
| --- | --- | --- | --- | --- | --- | --- |
| **2** | -0.42±0.92 | 0.49±0.01*** |  |  | 0.85 | 200 |
| **3** | -1.51±1.01 | 0.48±0.02*** | 0.17±0.08* |  | 0.83 | 196 |
| **3** | -1.66±1.41 | 0.48±0.02*** |  | 0.05±0.04 | 0.83 | 191 |
| **4** | -1.14±1.43 | 0.48±0.02*** | 0.18±0.10 | -0.01±0.05 | 0.83 | 190 |

Regressions used the validation data set of extracted Chl from 2022-2023 (cf. Fig. 5) and the concurrent measurements on the same samples using the Aquafluor (IVChl and CDOM) and Aquafast (Turbidity). Results are the fitted coefficients ± the standard error. The highest R^2^, adjusted for the number of variables, is using only IVChl as the independent variable.

*Fitted coefficient significant at *p*<0.05

***Fitted coefficient significant at *p*<<0.001
